# Genomic loci involved in sensing environmental cues and metabolism affect seasonal coat shedding in *Bos taurus* and *Bos indicus* cattle

**DOI:** 10.1101/2022.12.14.520472

**Authors:** Harly J. Durbin, Helen Yampara, Troy N. Rowan, Robert D. Schnabel, James E. Koltes, Jeremy G. Powell, Jared E. Decker

## Abstract

Seasonal shedding of winter hair at the start of summer is well studied in wild and domesticated populations. However, the genetic influences on this trait and their interactions are poorly understood. We use data from 13,364 cattle with 36,899 repeated phenotypes to investigate the relationship between hair shedding and environmental variables, single nucleotide polymorphisms, and their interactions to understand quantitative differences in seasonal shedding. Using deregressed estimated breeding values from a repeated records model in a genome-wide association analysis (GWAA) and meta-analysis of year-specific GWAA gave remarkably similar results.

These GWAA identified hundreds of variants associated with seasonal hair shedding. There were especially strong associations on chromosomes 5 and 23. Genotype-by-environment interaction GWAA identified 1,040 day length-by-genotype interaction associations and 17 apparent temperature-by-genotype interaction associations with hair shedding, highlighting the importance of day length on hair shedding. Accurate genomic predictions of hair shedding were created for the entire dataset, Angus, Hereford, Brangus, and multi-breed datasets. Loci related to metabolism and light-sensing have a large influence on seasonal hair shedding. This is one of the largest genetic analyses of a phenological trait and provides insight for both agriculture production and basic science.

## Introduction

Most mammals replace their coat or molt either completely or incompletely at annual or bi-annual intervals as an adaptive response to seasonal and climatic variation (Beltran *et al*. 2018). In cattle, molting occurs annually in the late spring and early summer when thick winter coats are exchanged for shorter ones in preparation for warmer temperatures. Generally, the onset of seasonal shedding is driven by hormone cascades initiated by the hypothalamus–pituitary–gonadal axis in response to environmental cues such as hours of sunlight per day (day length) and changes in temperature (Helm *et al*. 2013). Among ungulates and other mammals, the effects of temperature and day length interact to induce seasonal molting (Gebbie *et al*. 1999; Zimova *et al*. 2014, 2018; Schmidt *et al*. 2017). This interaction has never been explicitly demonstrated in cattle, although Yeates (Yeates 1955) showed that artificial manipulation of day length can be used to perturb the timing of hair coat shedding regardless of temperature, while Murray (Murray 1965) found a moderate effect of temperature on hair coat shedding among cattle at similar latitudes.

The timing and completeness of molting is also influenced by variables intrinsic to the individual, including plane of nutrition, life stage, and social status (Cowan *et al*. 1972; Heydon *et al*. 1995; Déry *et al*. 2019). In some species, inaccurate molt timing has a high fitness cost, and therefore phenotypic plasticity is limited (Zimova *et al*. 2014). In other species (including cattle), variation in molting has been documented within groups of contemporary individuals (Turner and Schleger 1960; Turner 1964), suggesting genetic variation influences an animal’s ability to respond to environmental cues. Despite the extensive body of research exploring its biological basis in wild populations, domestic populations, and humans, very few studies have focused on the genetic basis of seasonal coat change (see (Ferreira *et al*. 2017, 2020)) and to our knowledge only Durbin et al., 2020 (Durbin *et al*. 2020) has associated genetic variants with phenotypic variation in coat change. Though the cost of a “mismatched” seasonal phenotype may be lower in domestic species like cattle compared to many wild populations, they still have large impacts on productivity (St-Pierre *et al*. 2003; Baumgard *et al*. 2015). Previous work has demonstrated the impact of poor early summer hair shedding upon economically relevant traits such as growth and milk production in cattle (Gray *et al*. 2011; Durbin *et al*. 2020). Here, we explore the genetic, environmental, and interaction effects that influence early summer hair shedding. We use a multi-breed, repeated records dataset of early summer hair shedding scores collected across a range of latitudes, environmental conditions, and production systems to investigate how light, temperature, and metabolism interact with genomic loci to affect the timing of summer hair shedding in cattle.

## Materials and methods

All analyses were reproducibly pipelined using Snakemake workflows (Köster and Rahmann 2018) and R (R Core Team 2020). Unless otherwise specified, the {tidyverse} (Wickham *et al*. 2019) suite of packages was used for data processing. Code is available at https://github.com/harlydurbin/mizzou_hairshed_public.

### Phenotypes

Hair shedding scores were collected over 9 years by 77 beef cattle producers and university groups, with most scores collected between 2016 and 2019 (Figure S1a). Hair shedding was classified on an integer 1-5 scale based on the systems developed by Gray et al. (2011) (Gray *et al*. 2011) and Turner & Schleger (1964) (Turner and Schleger 1960) as described in Durbin et al. (2020) (Durbin *et al*. 2020), where a score of 5 indicated 100% winter coat remaining and a score of 1 indicated 0% winter coat remaining. Participants were asked to hair shedding score cattle when they observed the greatest amount of variation in shedding between contemporary individuals. Most herds were hair shedding scored once per year between mid-April and mid-June, but some groups chose to score cattle multiple times across the span of several months.

This resulted in between 1 and 8 scores per animal per year. Most cattle were scored in at least two separate years (8,839 or 66.11% of all individuals; Figure S1b).

When an animal’s date of birth was available, its “age class” was calculated based on the date that the score was recorded. When no date of birth was available, the producer-provided integer age was used. Unreported score dates were assumed to be May 1 of the scoring year for the purposes of age class calculation. Age class was calculated as *(n*365d)-90d* to *(n+1)*365d-90d*, where *n* is the age classification and *d* is days. This means that animals at least nine months of age that had not yet reached their first birthday could still be classified as yearlings and so on. Age class calculations were based on the Beef Improvement Federation age-of-dam definitions (Cundiff *et al*. 2018) as in (Durbin *et al*. 2020), and hair shedding scores recorded on animals fewer than 275 (i.e., 365-90) days of age were excluded. Animals with differing sexes reported across multiple years were also excluded. Finally, hair shedding scores recorded on bulls and steers were excluded as they comprised < 5% of the data, and work in other species suggests the biological mechanisms underlying molting may be different between sexes (Déry *et al*. 2019). After filtering, 36,899 phenotypes from 13,364 cattle were retained for analysis.

### Genotypes & imputation

Genotypes from SNP arrays were available for 10,511 phenotyped individuals and an additional 1,049 relatives. These genotypes originated from multiple commercial and research assays varying in density from 26,504 to 777,962 markers. Most animals were genotyped with the GGP-F250, a research assay enriched for low-frequency and putatively functional SNPs (Rowan *et al*. 2019). On an assay-by-assay basis, markers with > 10% missing data and markers significantly deviating from Hardy-Weinberg equilibrium (p-value < 10^-50^) were set to missing. After marker-level filtration, samples with > 10% missing data were removed. The remaining genotypes for all assays were merged by position, with discordant calls set to missing for individuals genotyped with more than one assay. The merged genotypes were then imputed to the union of the GGP-F250 and Illumina BovineHD assays using a multi-breed reference panel and the two-step approach described in (Rowan *et al*. 2019). Finally, SNPs with a minor allele frequency below 1% were removed, resulting in genotypes at 747,009 markers for 11,560 individuals. All genotype quality control was conducted using PLINK (Purcell *et al*. 2007; Purcell 2020).

### Generation of the pedigree and genomic relatedness matrices

Using records provided by various participating breed associations, a three-generation pedigree was constructed for registered animals with at least one phenotype retained for analysis. This pedigree was then supplemented with parentage information provided by project participants for un-registered and commercial animals with a registered sire and/or dam. To increase pedigree connectedness, the American Angus Association registration number was used for cross-registered American Angus individuals, sires, and dams with records in more than one breed association.

Parentage was validated for genotyped animals with at least one genotyped parent using the SeekParentF90 program (Hayes 2011; Misztal *et al*. 2014). Based on imputation accuracy, the expected rate of genotyping error, and the distribution of Mendelian conflicts across all parent-progeny comparisons, parents found to have > 0.05% SNPs in Mendelian conflict with reported progeny were set to missing in the pedigree. In total, 106 sires and 130 dams were excluded for 236 individuals. The final three-generation pedigree consisted of 13,221 un-phenotyped relatives in addition to the 13,364 phenotyped animals, with 6,733 unique sires and 17,954 unique dams. In order to take advantage of information from both genotyped and un-genotyped individuals, this pedigree was blended with genomic data to create the “hybrid” relationship matrix inverse (**H^-1^**) (Aguilar *et al*. 2011). **H^-1^** is calculated as:

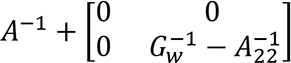

where **A^-1^** represents the inverse of the numerator relationship matrix, **A_22_^-1^** represents **A^-1^** subset to genotyped individuals, and **G^-1^** represents the inverse of the genomic relatedness matrix (GRM) calculated using the VanRaden method (VanRaden 2008). Genomic and pedigree relatedness matrices were blended with a weight of 0.05. In all models including a random effect of direct genetics, this matrix was used to represent relationships between individuals unless specified otherwise.

### Estimation of breeding values and genetic parameters

#### Full dataset

Genetic parameters and estimated breeding values (EBVs) of hair shedding were first estimated using records from all available animals in the following repeated records animal model, implemented with AIREMLF90 (Misztal *et al*. 2014).

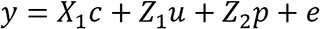

In this model, ***y*** represents a vector of hair shedding score phenotypes. Contemporary group effects are represented in the vector ***c***, where ***X_1_*** is a matrix relating the elements of ***c*** to ***y***. Contemporary groups were defined by the combination of herd ID, year, calving season (spring or fall), age group, toxic fescue grazing status, and score group. Based on the results of a model with age-in-years fit as a categorical fixed effect, age groups were defined as a) 1, b) 2-3, c) 4-9 or d) 10+. Grazing of tall fescue grass (*Lolium arundinaceum*) infected with the endophytic fungus *Epichloë coenophiala* has been shown to affect hair coat shedding in beef cattle (Gray *et al*. 2011; Durbin *et al*. 2020). Toxic fescue grazing status, or whether cattle grazed endophyte-infected fescue in spring of the recording year, was reported by participants as yes or no. Score group was used to account for differences in scoring dates within a herd and a year. In cases where an entire herd was not scored on the same day in a given year, records were assigned to a score group using a 5-day sliding window that maximized group size.

Records from contemporary groups with fewer than 5 records were discarded. Additional models with various effects fit separately from contemporary group or removed from the model were fit to test the effects of environment and management on hair shedding.

#### Breed-specific datasets

When performing single-step genomic best linear unbiased prediction (ssGBLUP) in crossbred populations, including data from both purebred and crossbred animals yields the highest prediction accuracy, assuming individuals have sufficiently similar genetic structure (Alvarenga *et al*. 2020). When individuals are not sufficiently similar, calculation of the GRM without accounting for differences in allele frequencies between populations can result in inflated estimates of inbreeding. In turn, this can cause inflated EBVs and associated reliabilities for some individuals. Further, animal models assume that all individuals in the pedigree derive from the same founder individuals. Violation of this assumption (as in the case of multi-breed and cross-bred evaluations) can result in inflated estimates of the additive genetic variance (Dong *et al*. 1988). Thus, we chose to replicate analyses completed in the full dataset in four breed-specific subsets with sufficient sample size for independent genetic evaluation.

Calculation of within-breed EBVs also allowed us to search for differences in additive genetic variation between breeds. The first through third datasets contained records from cattle registered with the American Angus Association (St. Joseph, MO; http://www.angus.org/), International Brangus Breeders Association (San Antonio, TX; https://gobrangus.com/), and American Hereford Association (Kansas City, MO; https://hereford.org/) respectively. The fourth dataset consisted of cattle registered with partner breed associations participating in the International Genetics Solutions (IGS) multi-breed evaluation (Bozeman, MT; https://www.internationalgeneticsolutions.com/). Further descriptions of these datasets can be found in Table 1. The three-generation pedigree and genotypes for associated animals were extracted for each dataset and used to construct the hybrid relatedness matrices **H^-1^** (see *Generation of the pedigree and relatedness matrices* section).

**Table 1.** Descriptions of the breed-specific datasets.

| Dataset | Description | N<br>genotyped<br>animals | N total<br>phenotypes |
| --- | --- | --- | --- |
| Angus | Purebred Angus cattle registered in the American Angus Association and commercial Angus cattle enrolled in the Breed Improvement Record Program | 3,286 | 8,674 |
| Brangus | Brangus and Ultrablack cattle registered in the International Brangus Breeders Association | 883 | 1,829 |
| Hereford | Purebred Hereford cattle registered in the American Hereford Association | 1,005 | 2,857 |
| IGS | Purebred and crossbred cattle registered in one or more of the breed associations participating in the International Genetics Solutions multi-breed evaluation: American Simmental Association, Red Angus Association of America, American Shorthorn Association, American Gelbvieh Association | 4,722 | 10,996 |

#### Evaluation of breeding values

Prediction models were evaluated in the full and breed-specific datasets using several metrics proposed by Legarra and Reverter (2018) and explored further by Macedo et al. (2020) (Legarra and Reverter 2018; Macedo *et al*. 2020). First, breeding values were estimated in ten separate iterations within each dataset, excluding phenotypes from a randomly selected 25% of individuals. Within each iteration, “partial” EBVs (*μ_p_*) were regressed on their corresponding “whole” EBVs (*μ_w_*) from the model including all animals. In the absence of dispersion, the resulting slope is expected to be 1. Next, we estimated bias (*Δ_p_*) by subtracting the absolute value of *μ_p_* from the absolute value of *μ*. Finally, prediction accuracies were calculated as *acc* 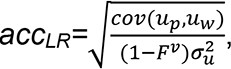 *cov(μ_p,_ μ_w_)* represents the covariance between partial and whole EBVs, *F^v^* represents the average inbreeding coefficient among animals with phenotypes randomly excluded, and *σ^2^_u_* represents the additive genetic variance of hair shedding score estimated using all individuals.

In order to evaluate the effect of explicitly accounting for breed structure on prediction accuracy and model fit, we tested a model including principal components from a PCA of all genotyped animals as fixed effects. The goal of this analysis was to determine if downstream analyses should use results from the previous (“full”) model discussed above. Principal component analysis was conducted using all 11,560 individuals and 747,009 SNPs with EIGENSOFT smartPCA v.7.2.1 (Patterson *et al*. 2006).

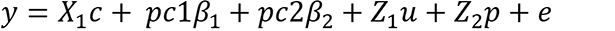

Only phenotypes from genotyped animals were included in this model. Otherwise, it was identical to the full model besides the inclusion of principal components 1 and 2 as covariates. This was compared to a model identical to the full model, except that only phenotypes from genotyped animals were included.

### The effects of temperature and photoperiod

Latitude and longitude coordinates were determined for each herd location using producer-provided addresses and the R package {tidygeocoder} (Cambon 2020). Based on these coordinates, the daily apparent high temperature, the sunrise time, and the sunset time were retrieved for the 30 days prior to each hair shedding scoring date at each unique geographic location using the {darksky} R package (Rudis 2017). The

{darksky} package interfaces with the Apple Dark Sky API to query NOAA historical weather records. For each score date-geographic coordinate combination, the resulting 30-day range of apparent high temperatures was then averaged. Apparent temperature can be thought of as a proxy for heat stress, as it combines the effects of real temperature, relative humidity, and wind speed. Similarly, day lengths were calculated by subtracting the time of sunrise from the time of sunset, then averaged across the 30- day range to act as a proxy for light exposure prior to hair scoring. Next, we fit the following repeated records animal model using AIREMLF90 (Misztal *et al*. 2014).

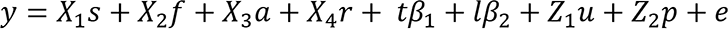

In this model, **y** is a vector of hair shedding score phenotypes; **s**, **f**, **a** and **r**, represent vectors of calving season, toxic fescue grazing status, age group, and year effects with matrices **X_1_**, **X_2_**, **X_3_**, and **X_4_** relating observations to effects; β_1_represents the regression of **y** on mean apparent high temperature (**t**) and β_2_ represents the regression of **y** on mean day length (**l**). The effect of farm or herd is confounded with the effect of latitude and by extension, both temperature and day length. Therefore, no herd effect was included.

Three additional models were also tested that were nearly identical to the base model above except for their inclusion of the temperature or day length variables. In two reduced models, only mean apparent high temperature or only mean day length were fit. In one expanded model, both temperature and day length were included plus an interaction effect, which was calculated by centering the individual variables then taking their product. All three of these models were compared to the base model using AIC and a likelihood ratio test.

#### Recommendations for genetic evaluations

In routine genetic prediction, additive and environmental variances are often partitioned by fitting a single contemporary group effect. For some traits, fitting an additional effect external to the contemporary group definition results in more accurate predictions despite the increased computational cost (i.e., the effect of age-of-dam fit for many maternally-influenced traits; (Cundiff *et al*. 2018)). For the purposes of large-scale genetic evaluations, it is of interest to know if the inclusion of additional environmental information provides a better fit than a simpler model including only contemporary group effects. To test this, we compared the base model and four repeated records animal models similar to those explored in the previous section. The first of these can be described as:

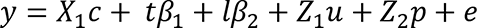

In this model, **y** represents a vector of hair shedding score phenotypes, and **c** represents contemporary groups defined in the same way as the model discussed in *Estimation of breeding values and genetic parameters*. Identical to the models fit in the previous section, β_1_represents the regression of **y** on mean apparent high temperature (**t**) and β_2_ represents the regression of **y** on mean day length (**l**). Two other models included only temperature or only day length alongside the contemporary group effect. The final, expanded model included an interaction between day length and temperature. In all models, records from contemporary groups smaller than 5 animals were removed.

### Genome-wide association

#### Deregression of breeding values and single-SNP regression

EBVs are an appealing pseudo-phenotype for further association studies as they represent the estimated additive genetic merit of an individual with environmental variance removed and combine repeated records into a single value. However, failing to account for the heterogeneous variances between EBVs resulting from the influence of familial data and in the case of repeated records traits, differing numbers of phenotypes per individual, can result in decreased power and increased false positive rate (Ekine *et al*. 2014). To take advantage of our repeated records, genome-wide association analyses (GWAA) were performed using deregressed breeding values (DEBVs).

First, reliabilities for EBVs were calculated as 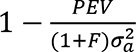 (Aguilar *et al*. 2020), where *PEV* represents the approximated prediction error variance and *F* represents the pedigree-based inbreeding coefficient for the animal of interest calculated using the R package {optiSel} (Wellmann 2020). Next, EBVs for genotyped animals were deregressed using the method proposed by Garrick et al. (2009) (Garrick *et al*. 2009), implemented in the {DRP} R package (Lopes 2016). The resulting 11,560 DEBVs were used as pseudo-phenotypes in SNP1101 single-SNP regression (Sargolazei 2014).

DEBVs were weighted by *(1/rel)-1*, where *rel* is the DEBV’s associated reliability with parent information removed as calculated using the Garrick et al., 2009 (Garrick *et al*. 2009) method. These weights were used to construct the **R^-1^** matrix of the mixed-model equations. Covariance between records due to relatedness was accounted for with a GRM constructed using the VanRaden method (VanRaden 2008). Post-analysis, p-values for single-SNP associations were adjusted by the estimated inflation factor (1.40) using the genomic control method (Devlin and Roeder 1999). After genomic control, p-values were converted to false discovery rate (FDR) adjusted q-values using the R package {qvalue} (Storey *et al*. 2017).

#### GWAA meta-analysis

Meta-analysis of summary statistics from multiple independent GWAA increases power and decreases false discovery rate compared with single-study results. In addition to GWAA of DEBVs, we tested meta-analysis of year-specific GWAA.

First, we fit 4 separate GWAA in GCTA-MLMA (Yang *et al*. 2011, 2014) using phenotypes recorded in each of the years between 2016 and 2019. Phenotypes were adjusted for environment and management using contemporary group designations as implemented in the *Estimation of breeding values and genetic parameters* section. In cases of animals with multiple phenotypes available within a given year, the phenotype recorded closest to May 1 was used. Records from contemporary groups with fewer than 5 records were excluded, resulting in 5,146, 4,989, 6,896, and 5,292 phenotypes included in the analyses associated with 2016, 2017, 2018, and 2019, respectively. Next, meta-analysis of the 4 year-by-year GWAA was performed using the first random effects (RE1) model implemented in METASOFT (Han and Eskin 2011). The RE1 method was used as it reported effect sizes and standard errors used in downstream analyses. The resulting meta-analysis p-values were adjusted by an inflation factor of 1.52 using the genomic control method (Devlin and Roeder 1999) and transformed to q-values (Storey *et al*. 2017).

#### Conditional & joint association analysis

Conditional and joint association analysis (COJO) is a stepwise model selection procedure used to identify independently associated SNPs and previously undetected secondary associations. Thus, it can be used to refine genome-wide association signals. COJO uses summary statistics (p-values as well as SNP effect betas and their associated standard errors) derived directly from a single GWAA or from meta-analysis of multiple GWAA. Betas and standard errors were not available from the analyses detailed in *Deregression of breeding values and single-SNP regression,* so we performed COJO using p-values obtained in the *GWAA meta-analysis* section.

Summary statistics from the METASOFT RE1 model were then used for COJO in GCTA (Yang *et al*. 2012) with linkage disequilibrium (LD) calculated within animals with a phenotype included in at least one of the year-by-year GWAA.

#### Genotype-by-environment interaction GWAA

Main effects GWAA is used to identify variants statistically associated with the phenotype of interest agnostic to the environment in which the phenotype was expressed, assuming appropriate model specification. By contrast, genotype-by-environment interaction (GxE) GWAA can be used to identify variants whose association with the phenotype of interest is dependent upon the environment in which the phenotype was expressed. Using the environmental data gathered in *The effects of temperature and photoperiod*, we fit GWAA modeling the interaction of genetics and either apparent high temperature or day length in GEMMA (Zhou and Stephens 2012). Phenotypes were pre-adjusted by subtracting the contemporary group fixed effect estimate (in this analysis defined as farm, year, calving season, age group, and fescue status) from the raw hair shedding score. As such, records were excluded from contemporary groups containing fewer than 5 animals. One GWAA was fit per environmental variable (mean apparent high temperature or mean day length) per year (2016-2019) and in cases of animals with multiple phenotypes per year, one phenotype was randomly chosen to increase variation in environmental data. Meta-analysis of the resulting summary statistics was then performed in METASOFT using the second random effects (RE2) model (Han and Eskin 2011). The ☐_mean_ (1.10 for temperature and 1.40 for day length) and ☐_hetero_ (0.47 for temperature and 0.33 for day length) calculated in this first run of METASOFT were used to adjust RE2 p-values in a subsequent run. Finally, adjusted RE2 p-values were transformed to q-values to correct for multiple testing (Storey *et al*. 2017).

#### Annotation and enrichment

Gene set and QTL enrichment analyses were performed separately for 1) the results of the main effect GWAA using DEBVs, 2) the main effect GWAA meta-analysis, 3) COJO selected SNPs, and 4) the GxE interaction effect GWAA meta-analysis of mean day length.

First, genes within 50 kb of variants significant at FDR < 5% were identified using the {GALLO} package (Fonseca *et al*. 2020) and ARS-UCD1.2 bovine genome coordinates (Rosen *et al*. 2020). In the case of COJO selected SNPs with no genes within 50 kb (n = 13), the nearest gene regardless of distance was included in the COJO gene set. Next, gene set enrichment analysis was performed for each gene set using the ‘gost’ function provided in the {gprofiler2} package (Kolberg and Raudvere 2020), which interfaces with the g:Profiler toolkit to query publicly available functional annotation databases. Considered data sources included gene ontology terms, KEGG pathways (Kanehisa and Goto 2000), Reactome pathways (Jassal *et al*. 2020), CORUM protein complexes (Giurgiu *et al*. 2019), and human disease phenotypes from the Human Phenotype Ontology database (Köhler *et al*. 2021). Significance values for gene set enrichment results were corrected for multiple testing using the g:SCS algorithm, which is designed for hierarchically related, non-independent tests.

We also performed QTL enrichment analysis using Animal QTLdb QTL annotations (Hu *et al*. 2019). The {GALLO} R package was used to identify QTL annotations within 50 kb of variants significant at FDR < 5% and to calculate enrichments. Significant enrichments were determined using a Benjamini-Hochberg adjusted p-value threshold of 0.05.

## Results

### Estimation of breeding values and genetic parameters

Models with and without principal components included as covariates were fit to assess the effect of population structure on breeding values. In principal component analysis, eigenvector 1 accounted for 35% of genetic variation, with Hereford animals at the negative end and Angus individuals at the other end. Eigenvector 2 accounted for differences in *Bos taurus* versus *Bos indicus* ancestry and explained 25% of the variance (Figure S2). The remaining principal components explained much less variation (Figure S3), and therefore we chose to include eigenvectors for only the first two principal components as covariates in the model evaluating the effect of explicitly accounting for breed structure. A likelihood ratio test comparing the models with and without PCs fit as fixed effects indicated that inclusion provided a moderately better fit (*- log_10_(p-value)* = 3.04). However, this model required the exclusion of phenotypes from un-genotyped animals. Further, including PCs resulted in lower prediction accuracy (*acc_LR_*) for hair shedding across 10 iterations (mean *acc_LR_* = 0.62 with PCs versus mean *acc_LR_* = 0.68 without PCs). Therefore, we chose to consider results from the “full” model excluding PCs and including all available individuals for downstream analyses.

Across the full and breed-specific datasets, narrow-sense heritability (*h^2^*) and repeatability (*r*) were similar to parameters reported for American Angus cattle by Durbin et al., 2020 (Durbin *et al*. 2020) and by Gray et al., 2011 (Gray *et al*. 2011) (Table 2). Estimated *h^2^* ranged from 0.32 (Hereford) to 0.41 (IGS) and estimated *r* ranged from 0.40 (Brangus, Hereford) to 0.48 (IGS). The additive genetic variance (*σ^2^_A_*) estimate in the full dataset fell within the range of estimates of *σ^2^_A_* in the breed-specific datasets, suggesting that this value was not inflated by the inclusion of crossbred animals. The permanent environmental variance (*σ^2^_PE_*) accounted for 5-7% of total variance with the exception of the Brangus dataset, for which *σ^2^_PE_* was essentially zero. The International Brangus Breeders Association did not begin participating in the project until 2018, and so 96% of Brangus animals had only one or two years of data. This likely explains why no permanent environmental effect was estimated in the Brangus dataset.

**Table 2.** Additive genetic, permanent environmental, and residual variance estimates as well as narrow-sense heritability and repeatability within the full dataset and each of the four breed-specific datasets. Approximated standard errors for heritability and repeatability estimates are in parentheses.

| **Dataset** | $\sigma^2_A$ | $\sigma^2_{PE}$ | $\sigma^2_E$ | $h^2$ | $r$ |
| --- | --- | --- | --- | --- | --- |
| Angus | 0.358 | 0.052 | 0.557 | 0.370<br>(0.022) | 0.424<br>(0.014) |
| Brangus | 0.267 | 0.000 | 0.403 | 0.400<br>(0.031) | 0.400<br>(0.031) |
| Hereford | 0.215 | 0.054 | 0.400 | 0.321<br>(0.044) | 0.402<br>(0.024) |
| IGS | 0.390 | 0.070 | 0.500 | 0.407<br>(0.020) | 0.480<br>(0.012) |
| Full dataset | 0.331 | 0.072 | 0.500 | 0.368<br>(0.012) | 0.448<br>(0.008) |

In the full dataset, the median EBV was -0.02, ranging from -2.32 to 1.92.

Though variation in EBVs largely overlapped between breeds, breeds recently selected for performance in the “show ring” where fuller hair coats are desirable (Shorthorn and Maine-Anjou) tended to have higher (i.e., less desirable) EBVs (Figure S4). Further, breeds with known *Bos indicus* ancestry (Brangus and Charolais; (Decker *et al*. 2014)) tended to have lower EBVs.

Estimated *acc_LR_* ranged from 0.657 to 0.674 across 10 iterations using all available data. Among the breed-specific datasets, *acc_LR_* tended to be lowest in the Hereford dataset and highest in the IGS dataset (Table S1). The mean *b^v^_w,p_* was between 1.00 and 1.04 in all datasets, suggesting minimal dispersion.

### The effects of temperature and photoperiod

Mean hours of sunlight per day ranged from 10.89 to 15.41 hours, averaging hours with a standard deviation of 0.74. The mean apparent high temperature ranged from 4.23 to 39.33 °C with a mean of 25.87 and standard deviation of 5.37. The base model including apparent temperature and day length provided a better fit over both the model with only apparent temperature (*-log_10_(p-value)* = 92.35) or day length (*- log_10_(p-value)* = 156.52), while the model including the interaction effect provided a better fit than the base model (*-log_10_(p-value)* = 3.42). The interaction model also had a lower AIC value than the base model and both of the reduced models (Table 3). The day length BLUE from this expanded model suggested that, on average, hair shedding score decreases by 0.45 units for each hour increase in the mean hours of sunlight in the 30 days prior to scoring hair shedding. Further, hair shedding score was predicted to decrease by 0.07 units with every 1°C increase in the mean apparent high temperature for the 30 days prior to scoring.

**Table 3.** BLUEs and AIC values from four increasingly complex models quantifying the effects of day length and temperature on hair shedding.

| Model | Day length BLUE | Temperature BLUE | Day length*temperature BLUE | AIC |
| --- | --- | --- | --- | --- |
| Day length + covariates | -0.830 (0.008) | - | - | 94972.420 |
| Temperature + covariates | - | -0.121 (0.001) | - | 94380.790 |
| Day length + temperature + covariates (base model) | -0.394 (0.013) | -0.074 (0.002) | - | 93537.380 |
| Day length + temperature + day length*temperature + covariates | -0.446 (0.016) | -0.072 (0.002) | -0.006 (0.001) | 93509.710 |

Calving season, toxic fescue grazing status, and age group BLUEs from all four models are in Table S2. In general, BLUEs for grazing toxic fescue tended to be higher than BLUEs for not grazing toxic fescue and BLUEs for spring calving tended to be higher than BLUEs for fall calving, congruent with trends reported by Durbin et al., 2020 (Durbin *et al*. 2020). The magnitude of the effect of grazing toxic fescue was estimated to be ∼ 4 times larger in the day length only model than in the temperature only model, and ∼ 2 times larger than in the models including both day length and temperature.

### Recommendations for genetic evaluations

Based on a series of likelihood ratio tests and AIC comparisons, the base model including temperature and day length without an interaction effect provided the best fit to the data. However, the direction of the signs changed from negative to positive for all temperature and day length BLUEs relative to the models without a contemporary group effect. Besides being incongruent with biological expectation, this is likely a sign of collinearity and suggests that including temperature or day length is redundant when contemporary groups are properly constructed. The combination of score group and farm ID captures these environmental conditions, and therefore we recommend that producers hair shedding score their entire herd on the same day in adequately sized score groups as are represented in our analyzed data.

### Genome-wide association

In the main effects GWAA using DEBVs, 377 variants were genome-wide significant at FDR = 5% (Figure 2a). Of the 413 variants passing a FDR = 5% level in the meta-analysis of year-specific main effects GWAA, 20 were selected by COJO, plus an additional 37 variants with FDR < 5% (Figure 2b). The 57 SNPs selected by COJO resided in 30 independent associations, with independent associations delimited by pairs of adjacent SNPs greater than 1 Mb apart as in (Yang *et al*. 2012).

**Figure 1.**
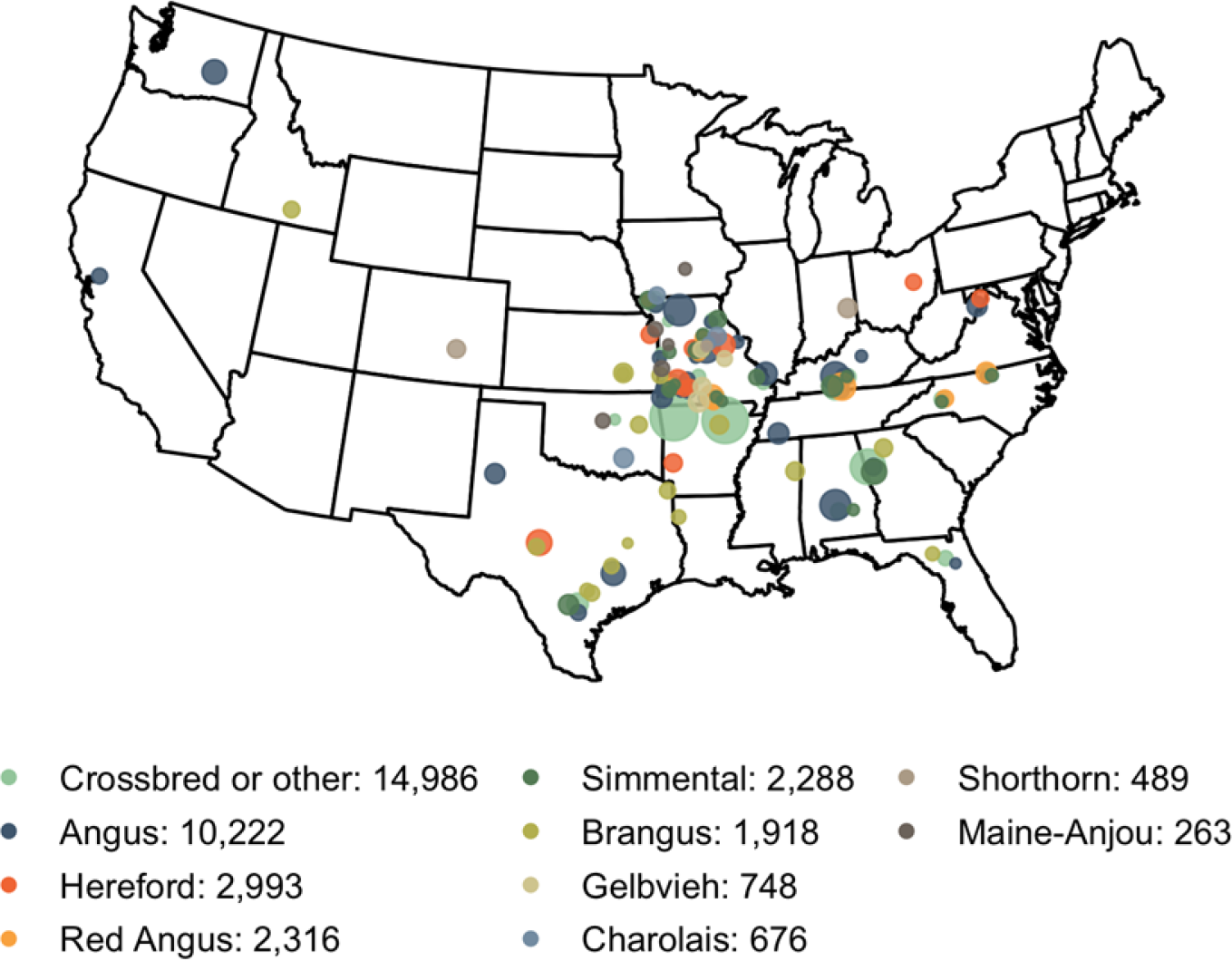
Counts of hair shedding score records by reported breed. Most phenotypes came from three breeds and were recorded in the Midwest or South. For the purposes of this map, Angus, Hereford, Red Angus, Simmental, and Gelbvieh animals with at least ⅝ ancestry assigned to the given breed based on pedigree estimates were included in that breed. Animals with unknown ancestry, less than ⅝ ancestry assigned to one breed, or of a breed not listed above were called “Crossbred or other”.

**Figure 2.**
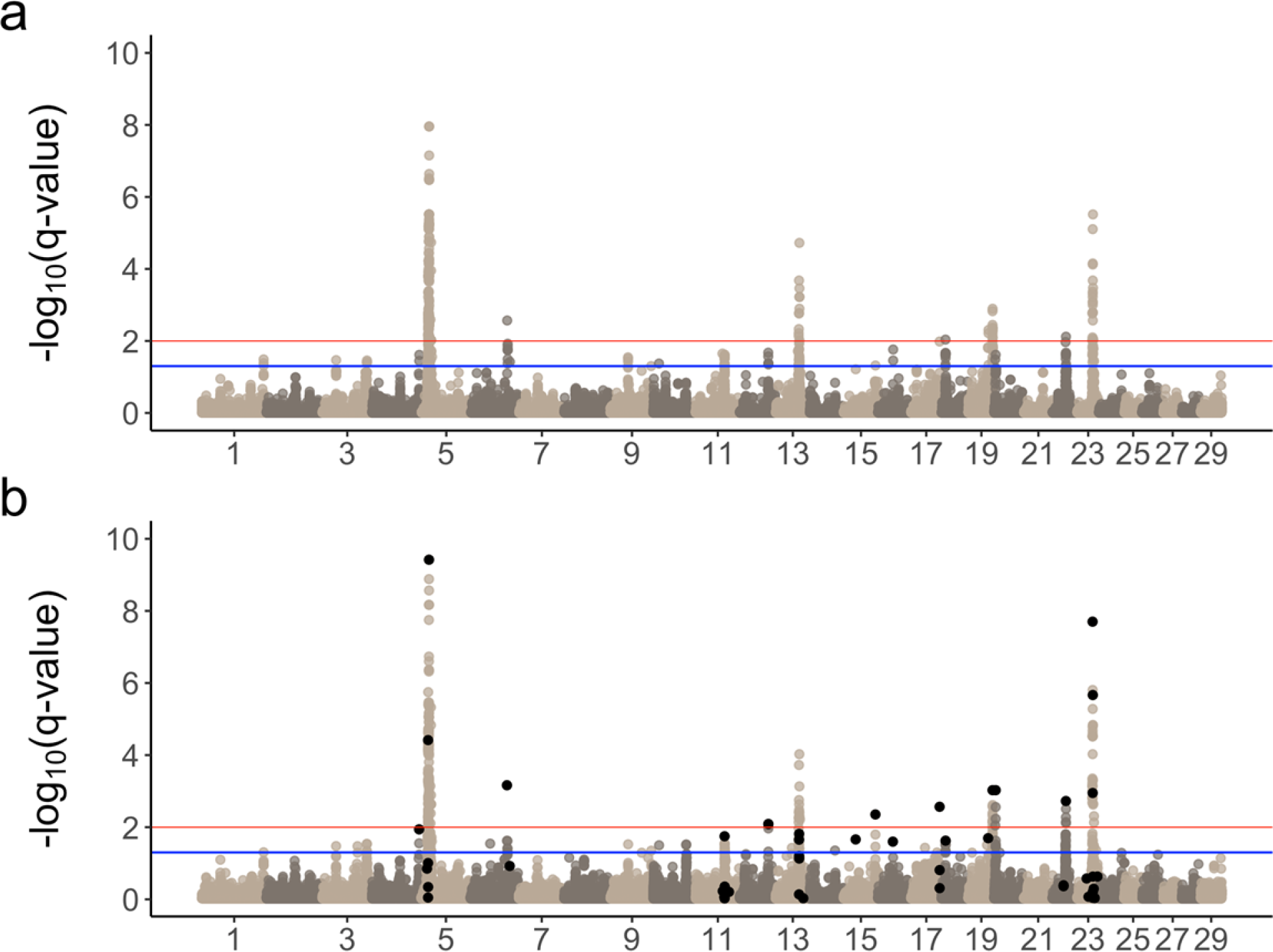
Manhattan plot of main effect *-log_10_(q-values)* **(a)** using DEBVs and **(b)** using meta-analysis. Blue lines represent FDR = 5% and red lines represent FDR = 1%. Black points represent variants identified by COJO selection.

Over half of all significant SNPs were found on chromosome 5 as in (Durbin *et al*. 2020) (Figure 3a). COJO selection suggests that these results might represent 3 separate causal loci. The lead SNP in both main effects GWAA (BTA5:18,767,155) was also the most significant SNP identified in COJO selection. The nearest gene to this variant is a lncRNA approximately 300 kb downstream (ENSEMBL ID *ENSBTAG00000053947*), followed by *KITLG* approximately 400 kb upstream.

**Figure 3.**
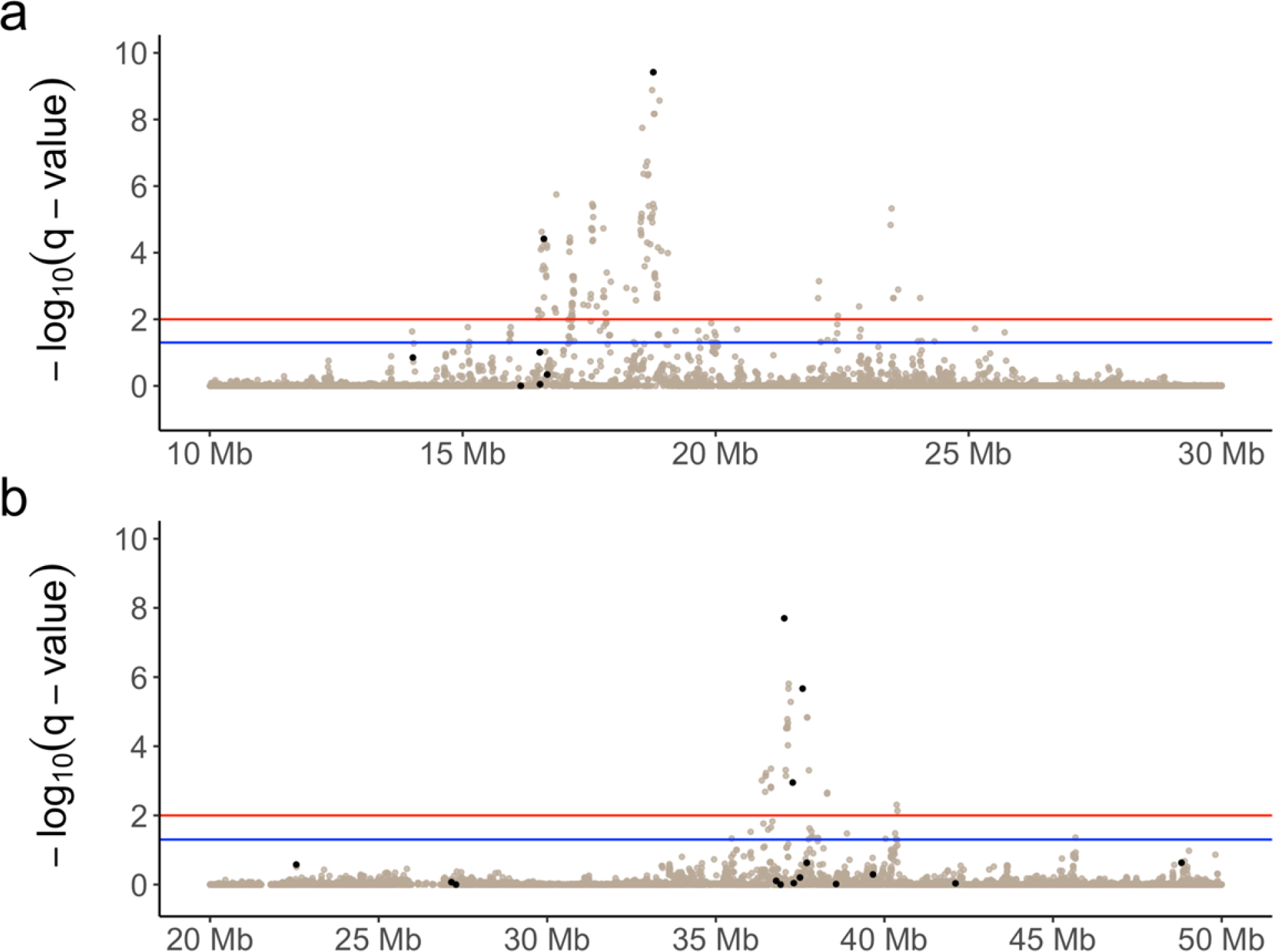
Manhattan plot of main effect *-log_10_(q-values)* on **(a)** chromosome 5, truncated from 10 Mb to 30 Mb, and **(b)** on chromosome 23, truncated from 20 Mb to 50 Mb. Blue lines represent FDR = 5% and red lines represent FDR = 1%. Black points represent all variants identified by COJO selection on chromosomes 5 and 23.

**Figure 4.**
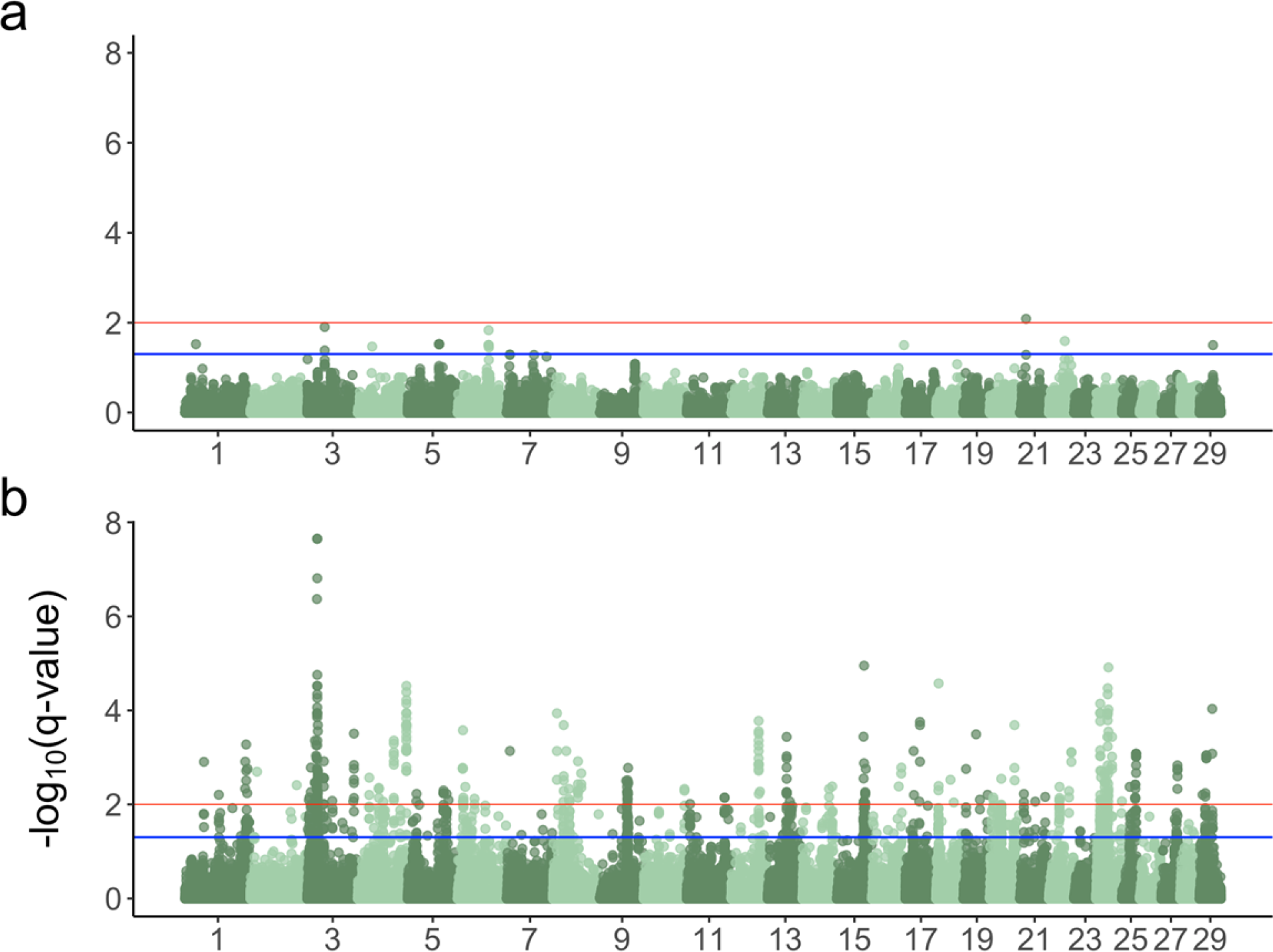
Manhattan plots of **(a)** mean apparent temperature and **(b)** mean day length interaction effect meta-analysis *-log_10_(q-values)*. Blue lines represent FDR = 5% and red lines represent FDR = 1%.

Gene set enrichment results were largely consistent between the two main effects GWAA strategies, both returning terms associated with regulation of apoptosis (Table S3). QTL enrichment analysis of the main effects results returned 6 significant terms (Table S2). Of these, “white spotting” was the most significantly enriched QTL term (p-value = 1.15 x 10^-16^). Upon further examination, this signal appeared to be driven by SNPs within and near to the *MITF* (*microphthalmia-associated transcription factor*) gene on chromosome 22, a master regulator of melanocyte production that is highly conserved across vertebrates (Steingrímsson *et al*. 2004; Levy *et al*. 2006; Hou and Pavan 2008).

Meta-analysis of mean apparent high temperature and mean day length GxE interaction effect GWAA resulted in 17 and 1,040 variants passing at FDR = 5%, respectively (Figure 3). Unlike main effect gene set enrichment results, most terms enriched in the analysis of day length GxE interaction results were associated with keratinization and cytoskeleton formation (Table S3).

## Discussion

In cattle, hair shedding is affected by nutrition and metabolism, temperature, and seasonal changes in the amount of daylight.

Calving season, toxic fescue grazing status, and age group BLUEs shed light on the relationship between seasonal hair shedding and nutrition/management effects. For example, younger cows have higher nutritional requirements and shed off later in the spring. Surprisingly, BLUEs for the oldest age group (cows aged 10 and up) were considerably more negative than BLUEs for mature cows (aged 4-9). In other mammals, patterns of seasonal coat shedding are typically “U-shaped” with age, with senescing animals in the last stage of their life typically shedding later than their younger counterparts (i.e., (Déry *et al*. 2019)). Since beef cows are typically culled from the herd or die between ages 11 and 12 (Azzam *et al*. 1990), it’s likely that the advantage predicted here for very old cows is reflective of selection (i.e., culling) allowing well-adapted individuals to remain productive later into their lives.

The estimated effect of grazing toxic fescue was largest in the day length only model. This could reflect the relationship between temperature and tall fescue ergot alkaloid levels (see (Klotz 2015) and the references therein), as major symptoms of fescue toxicosis is vasoconstriction increasing heat stress and decreased feed intake. Fescue toxins are also known to decrease serum prolactin (Klotz 2015), thus highlighting the relationship between fescue toxins, prolactin, and hair shedding.

Selection for more rapid hair shedding may not only increase heat tolerance, but may also identify cattle more resilient to fescue toxicosis.

The length of daylight hours has a tremendous impact on hair shedding. The hormonal cascades underlying seasonal coat change, largely driven by prolactin, are highly conserved between species. In response to changes in day length, photoperiodic cues are forwarded from the eye to the pineal gland, which is responsible for the production of melatonin. Melatonin inhibits prolactin, and decreased melatonin production from increased daylight allows for elevated serum prolactin, thereby triggering the spring molt (Zimova *et al*. 2018). Temperature changes have also been implicated via interaction with day length (Gebbie *et al*. 1999; Zimova *et al*. 2014; Schmidt *et al*. 2017), though the mechanisms by which temperature influences seasonal biology (and by which mammals sense temperature in general) are unclear (Caro *et al*. 2013). The day length-temperature interaction has never been explicitly demonstrated in cattle, though Peters and Tucker (Peters and Tucker 1978) showed the effect of photoperiod on serum prolactin is magnified by ambient temperature. Results from the models here quantifying the effects of external environmental variables on hair shedding score clearly point to roles for day length and temperature in regulating seasonal coat change. However, GxE GWAA of the two variables suggests a larger independent role for day length than temperature. Even though the two GxE GWAS had the exact same sample sizes, we observed substantially more associations in the daylength GxE GWAS. The lack of temperature GxE GWAA results found here is consistent with previous work demonstrating the inability of temperature alone to initiate seasonal coat change (Zimova *et al*. 2018). Our results show that in cattle, there is little to no genetic variation for temperature sensing. However, there appears to be a large number of genetic interactions with day length that are associated with hair shedding variation.

Using two very different GWAA approaches (DEBVs as pseudo-phenotypes and meta-analysis of year-specific GWAA), we found nearly identical results (Figure 2; Table S3; Table S4). In both main effects GWAA, over half of all significant variants were located on chromosome 5 (Figure 2). Durbin et al. (Durbin *et al*. 2020) found a similarly large association on chromosome 5 in American Angus cattle but was unable to narrow down candidate genes. They theorized that the strength of the association, likely caused by extensive long-range LD, could contain multiple causal mutations affecting multiple genes (Cannon and Mohlke 2018). Using COJO selection, we find evidence for three separate associations between 14 and 24 Mb on chromosome 5 (Figure 2b). The third association contains the lead COJO SNP and the lead SNP in both main effects GWAA, located at BTA5:18,767,155. The closest genes to this SNP are an unannotated lncRNA and *KITLG*, ∼ 300 kb downstream and ∼ 400 kb upstream, respectively. *KITLG* and its receptor, *KIT* have a multitude of roles across tissues, including in the retina.

Recently, *KITLG* was shown to protect against retinal degenerative diseases by preventing photoreceptor death in a mouse model (Li *et al*. 2020). Also of interest, *KIT* and *KITLG* regulate the activity of *MITF*, although the mechanism by which this happens is unclear (Hou and Pavan 2008). The gene *CEP290* is between the 2nd and 3rd COJO peaks, and its action is involved in the biogenesis of the photoreceptor sensory cilia (Rachel *et al*. 2012). Finally, *NTS* (neurotensin) is located at 15.5 Mb on chromosome 5; neurotensin is known to affect prolactin levels (McCann *et al*. 1982).

This connection to prolactin is interesting, based on the effect of prolactin on seasonal turnover of the haircoat.

The second largest association in the main effects GWAA was located on BTA23 in proximity to *PRL* (BTA23:35,332,705-35,341,607; Figure 3b), the gene encoding prolactin. As previously mentioned, prolactin is the main hormonal regulator of seasonal response, making it an obvious candidate gene. Further, mutations in *PRL* and its receptor were previously linked to an abnormal hair coat and thermoregulatory phenotype (Littlejohn *et al*. 2014). However, this association also contained *CDKAL1* (*CDK5 regulatory subunit associated protein 1 like 1*; BTA23:36,745,642-37,444,814), with many significant variants (including 5 COJO selected SNPs) within the gene itself (Figure 3b). In multiple human populations, *CDKAL1* variants are associated with risk for type 2 diabetes mellitus and decreased insulin secretion (Steinthorsdottir *et al*. 2007). These results could point towards a relationship between metabolic regulation and seasonal hair shedding in cattle. In support of this idea, mature body weight and 18-month body weight QTL associations were enriched in the main effects results (Table S4). Further, genes associated with “metabolism” (Table S3) and QTL associated with “metabolic body weight” and “ketosis” (Table S4) were enriched in the day length GxE GWAA results. Photoperiodic response is associated with patterns of seasonal growth and metabolic change across taxa in addition to coat change. When resources are more seasonally dependent, seasonal growth and nutrient partitioning is more extreme with higher adiposity during days with few hours of sunlight. Domestic animals are less dependent upon seasonal resources, but there is still evidence that growth, nutrient partitioning, and milk production respond to changing day length in livestock species (Petitclerc *et al*. 1983; Tucker *et al*. 1984; Barenton *et al*. 1988; Dahl *et al*. 2000; Small *et al*. 2003).

The hair follicle cycle is composed of three main phases highly conserved across mammalian species (Stenn and Paus 2001): anagen, catagen, and telogen. During catagen, the hair follicle involutes via apoptosis (Botchkareva *et al*. 2006), or the process by which cells are eliminated after fulfilling a biological purpose. Our gene set enrichment analyses of the main effects GWAA results returned multiple terms associated with regulation of apoptosis. Additionally, terms associated with keratin formation were enriched in the day length GxE gene set enrichment analysis. In a transcriptomic study of mountain hare seasonal coat color, Ferreira et al. (Ferreira *et al*. 2020) found similar enrichment of cell death terms and keratinization. This might suggest shared mechanisms between multiple forms of seasonal coat change.

QTL associated with “white spotting” were significantly enriched in the main effects results, driven by SNPs near and within *MITF* on BTA22. Perturbations to *MITF* are responsible for several audio-visual disorders across taxa, including Waardenburg syndrome (Tassabehji *et al*. 1994) and Tietzs syndrome (Amiel *et al*. 1998) in humans. Of particular interest, *MITF* is also essential for regulating the production of retinal pigment epithelial cells, which support the parts of the eye responsible for light sensing and color vision (Wen *et al*. 2016; García-Llorca *et al*. 2019; Han *et al*. 2020). In cattle and other mammals, light stimulation in the eye activates the pineal gland, which is the main regulator of photoperiodic responses including coat molting (Reiter 1991).

In the future, imputation of genotypes to sequence level would enable fine-mapping significantly associated regions and their functional annotations. At the genotype density used here, causal variants are unlikely to have been directly assayed. Sequence-level data in combination with our multi-breed dataset could enable the refinement of causal variants to near the base-pair level.

Data collected and maintained by non-professionals are often under-utilized, as it can introduce certain errors and biases. However, it can afford researchers an increased analytical power via vastly increased sample size when statistically modeled correctly. In a similar effort, Nowak et al. (Nowak *et al*. 2020) quantified the effects of temperature and day length on molting in mountain goats using data and photographs collected by non-professionals. Similarly, we were able to explore the functional biology of a complex trait using farmer-sourced data. This work reinforces the utility of “citizen-science” data collected by non-professionals as a powerful tool for studying complex trait biology.

## Conclusions

We confirm once again that hair shedding is moderately heritable with consistent estimates of heritability and repeatability between datasets. Using a crossbred and multi-breed dataset, we were able to show that a previously published association found on chromosome 5 in American Angus cattle is likely driven by multiple causal variants at this locus. Together, these results point towards important roles of daylight sensing and temperature in regulating bovine seasonal hair coat shedding and provide compelling candidate regions for functional analyses. Particularly, there appears to be a clear relationship between variation in hair shedding and ocular function. Despite a vast body of research exploring the biological mechanisms regulating seasonal molting across the tree of life, to our knowledge there have been no previous studies of how genetic loci contribute to individual variation in seasonal molting. Additionally, the photoperiodic and light-sensing mechanisms regulating most seasonal phenotypes, including coat shedding, is largely shared across species (see (Helm *et al*. 2013) and references therein). Therefore, this work also provides an important stepping off point for research in other species.

## Data Availability

Genotype, phenotype, and metadata are available from the Dryad Digital Repository (https://doi.org/10.5061/dryad.ngf1vhhz4).

## Acknowledgements

We thank the following participating breed associations for data, pedigree, and genotype sharing:

- The American Simmental Association, particularly Jackie Atkins, Rachel Endecott, and Jordan Bouwman
- The Red Angus Association of America, particularly Ryan Boldt
- The American Shorthorn Association, particularly Matt Woolfolk
- The American Gelbvieh Association, particularly Will Fiske and Taylor Buckley
- The American International Charolais Association, particularly Sally Northcutt
- The American Hereford Association, particularly Stacy Sanders
- The American Angus Association, particularly Steve Miller, Duc Lu, Kelli Retallick, and Dan Moser
- The International Brangus Breeders Association, particularly Macee Prause Thank you to all undergraduate student workers and interns who helped with data collection, organization, and accession of DNA samples from 2016-2019:
- Will Shaffer
- Natalie Barr
- Sarah Van Ausdal
- Brian Arisman
- Aaron Mott

Thank you to the farm staff at the Savoy and Batesville Research Stations with the Division of Agriculture at the University of Arkansas System, in particular Don Hubbell and John Tucker at the Livestock and Forestry Research Station who coordinated extensive hair score phenotyping for this study.

Thank you especially to the farmers and ranchers who contributed data, without whom these analyses would not have been possible.

## Funding

This project was supported by Agriculture and Food Research Initiative Competitive Grant no. 2016-68004-24827 from the USDA National Institute of Food and Agriculture.

## Conflict of Interest

Jared Decker and James Koltes are on the scientific advisory board of Vytelle, LLC.

## Supplementary File 1

**Figure S1.**
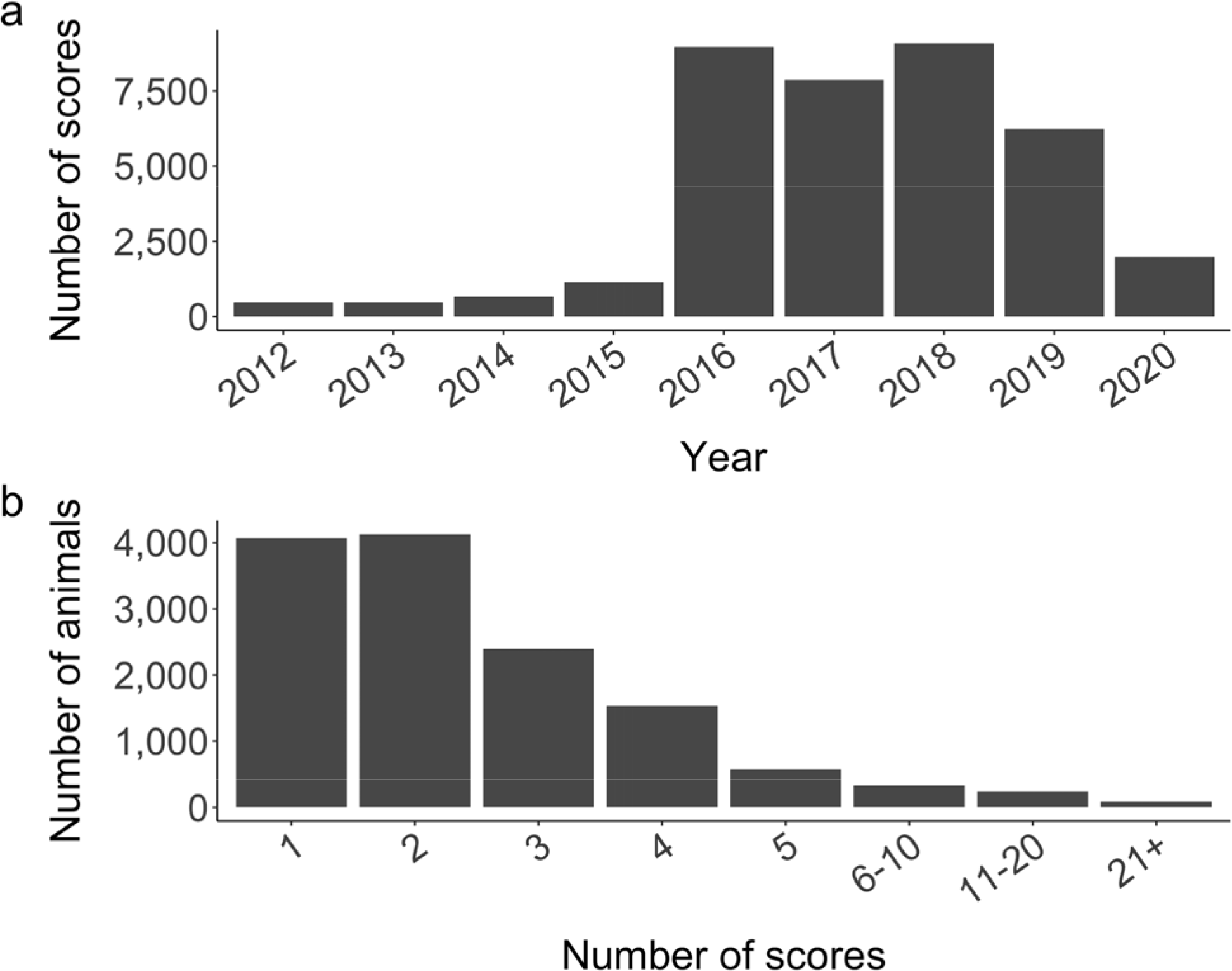
Counts of **(a)** hair shedding scores per year and **(b)** scores per animal across all years.

**Figure S2.**
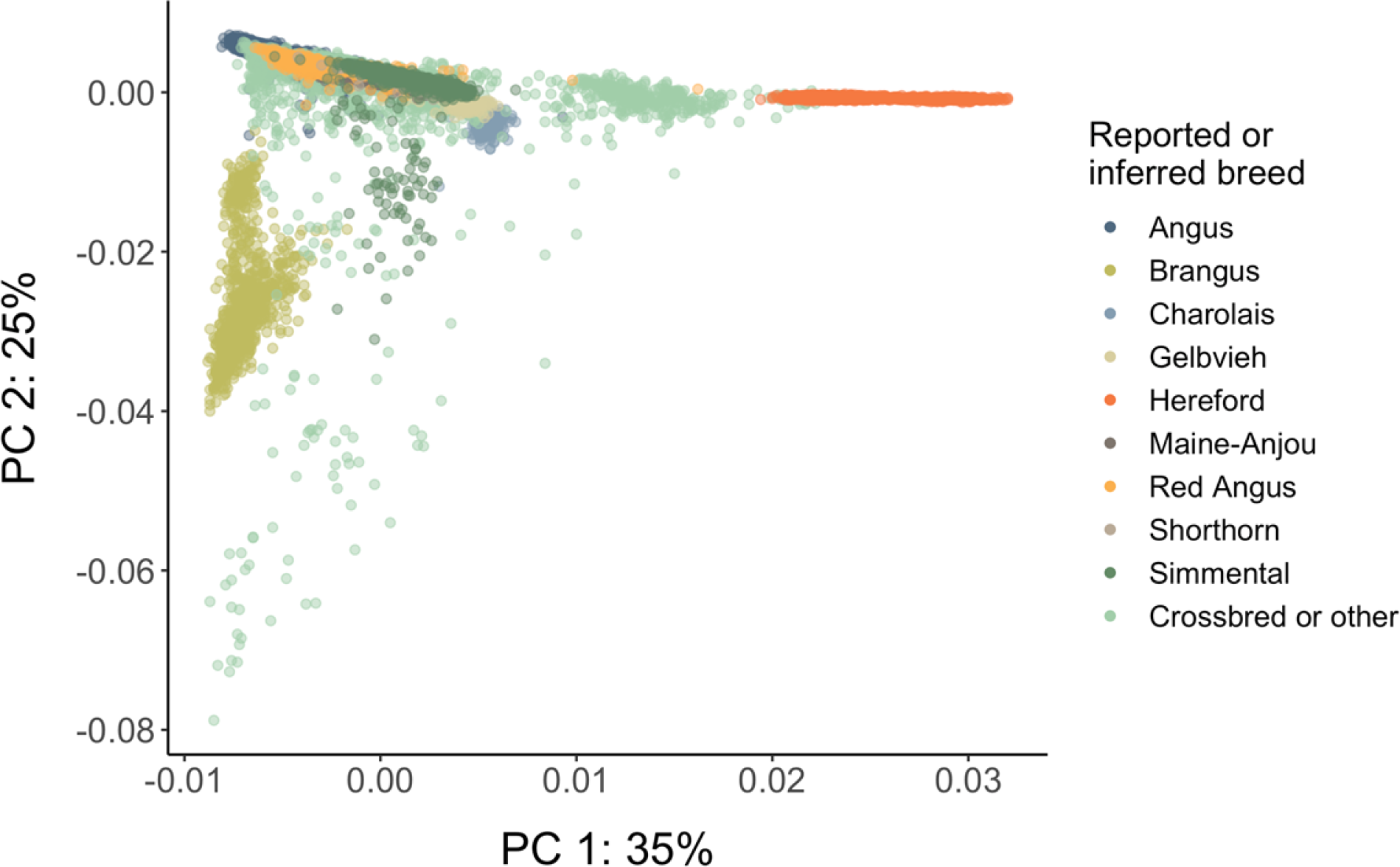
Principal components 1 and 2. For the purposes of this visualization, Angus, Hereford, Red Angus, Simmental, and Gelbvieh animals with at least ⅝ pedigree ancestry assigned to the given breed based on pedigree estimates were included in that breed. Animals with unknown ancestry, less than ⅝ pedigree ancestry assigned to one breed, or of a breed not listed above were called “Crossbred or other”.

**Figure S3.**
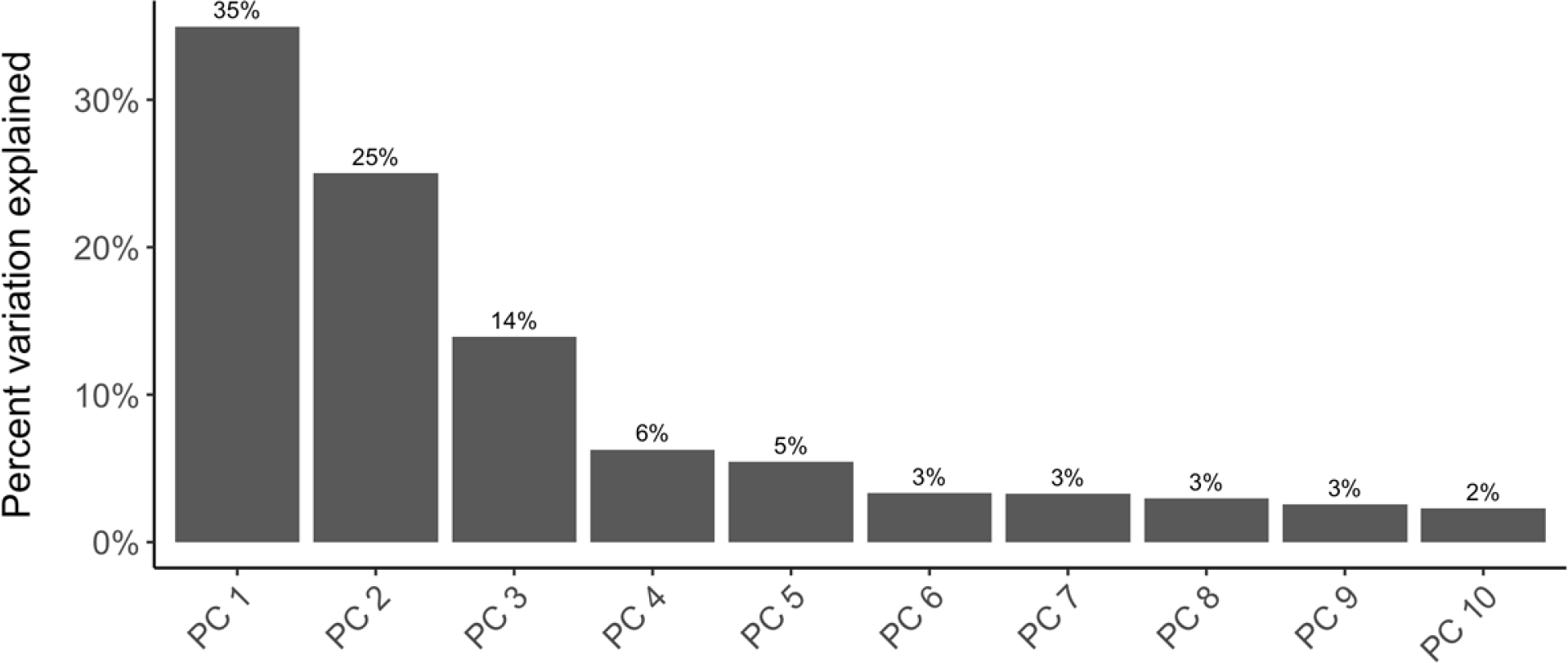
Principal components 1 and 2 explain 65% of variation in a principal component analysis of all 11,560 genotyped animals in the dataset.

**Figure S4.**
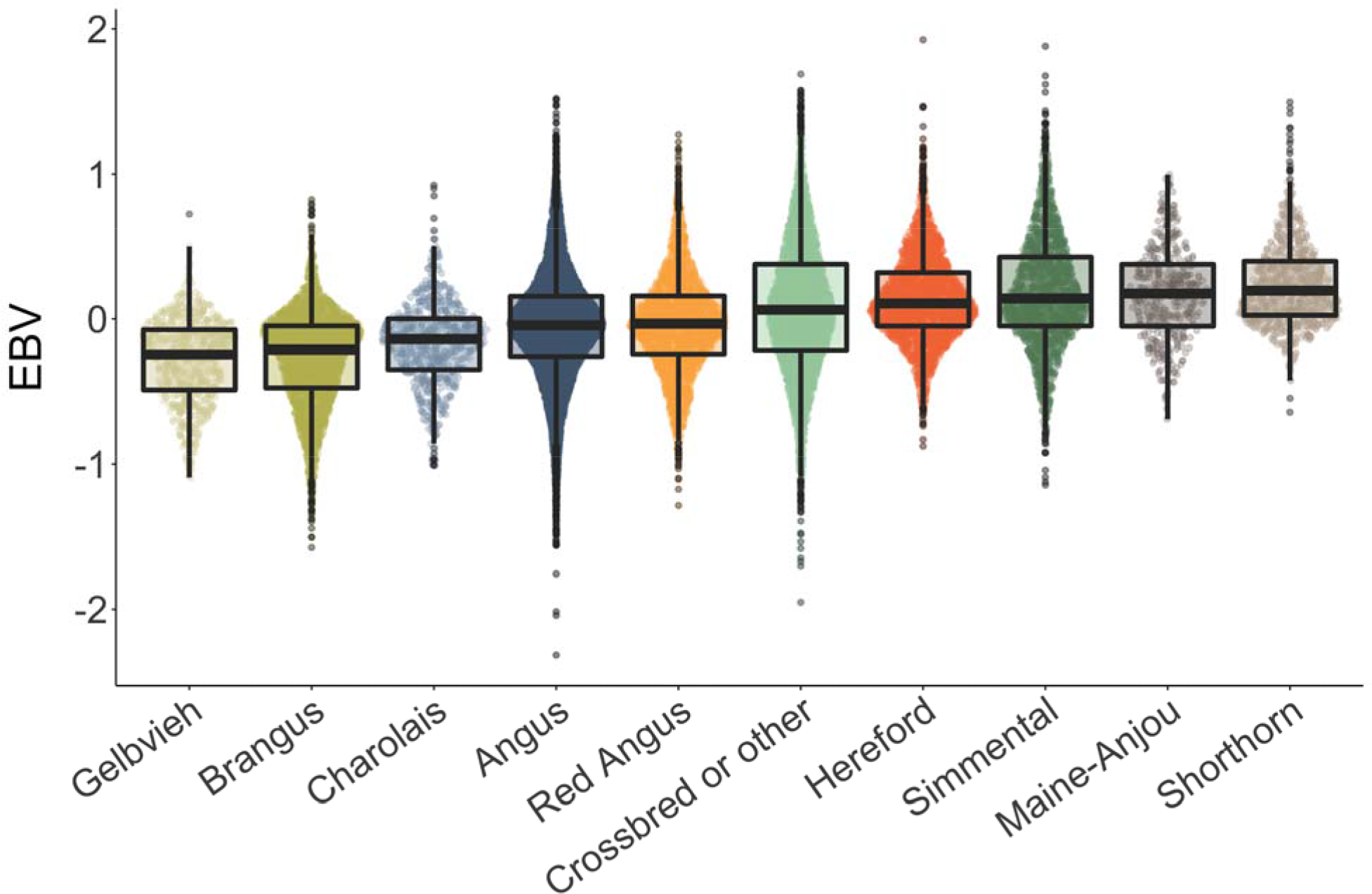
Comparison of EBVs from the full dataset by breed. For the purposes of this summarization, Angus, Hereford, Red Angus, Simmental, and Gelbvieh animals with at least ⅝ pedigree ancestry assigned to the given breed based on pedigree estimates were included in that breed. Animals with unknown ancestry, less than ⅝ pedigree ancestry assigned to one breed, or of a breed not listed above were called “Crossbred or other.”

**Table S1.** Evaluation of breeding values in the full and breed-specific datasets across 10 iterations.

| *Prediction accuracy* |  |  |  |  |
| --- | --- | --- | --- | --- |
| **Dataset** | **Minimum** | **Mean** | **Maximum** | **SD** |
| Angus | 0.585 | 0.594 | 0.601 | 0.006 |
| Brangus | 0.514 | 0.524 | 0.536 | 0.007 |
| Hereford | 0.498 | 0.520 | 0.545 | 0.013 |
| IGS | 0.653 | 0.663 | 0.676 | 0.007 |
| Full dataset | 0.657 | 0.665 | 0.674 | 0.006 |
| *Dispersion ($b_{w,p}^v$)* |  |  |  |  |
| **Dataset** | **Minimum** | **Mean** | **Maximum** | **SD** |
| Angus | 0.931 | 1.007 | 1.089 | 0.055 |
| Brangus | 0.903 | 1.014 | 1.116 | 0.072 |
| Hereford | 0.853 | 1.027 | 1.252 | 0.133 |
| IGS | 0.964 | 1.036 | 1.154 | 0.049 |
| Full dataset | 0.975 | 1.009 | 1.041 | 0.021 |
| *Bias ($\Delta_p$)* |  |  |  |  |
| **Dataset** | **Minimum** | **Mean** | **Maximum** | **SD** |
| Angus | 0.000 | 0.091 | 1.356 | 0.107 |
| Brangus | 0.000 | 0.069 | 0.964 | 0.091 |
| Hereford | 0.000 | 0.072 | 1.779 | 0.088 |
| IGS | 0.000 | 0.094 | 1.397 | 0.113 |
| Full dataset | 0.000 | 0.084 | 1.966 | 0.097 |

**Table S2.** BLUEs for toxic fescue grazing status across four increasingly complex models quantifying the effects of daily sunlight duration and temperature on hair shedding with approximated standard errors in parentheses. “Fall calving” BLUEs are relative to “spring calving” BLUE = 0, “not grazing toxic fescue” BLUEs are relative to “grazing toxic fescue” BLUE = 0, and age group BLUEs are relative to “age: yearling” BLUE = 0.

| Model | Fall calving | Not grazing toxic fescue | Age: 2-3 year olds | Age: 4-9 year olds | Age: 10+ |
| --- | --- | --- | --- | --- | --- |
| Day length + covariates | -0.122<br>(0.020) | -0.222<br>(0.022) | 0.091<br>(0.019) | -0.057<br>(0.021) | -0.206<br>(0.028) |
| Temperature + covariates | 0.001<br>(0.019) | -0.055<br>(0.022) | 0.106<br>(0.019) | -0.077<br>(0.021) | -0.231<br>(0.027) |
| Day length + temperature + covariates | -0.056<br>(0.019) | -0.124<br>(0.022) | 0.100<br>(0.018) | -0.067<br>(0.021) | -0.215<br>(0.027) |
| Day length + temperature + day length*temperature + covariates | -0.061<br>(0.019) | -0.136<br>(0.022) | 0.102<br>(0.018) | -0.061<br>(0.021) | -0.208<br>(0.027) |

**Table S3.**
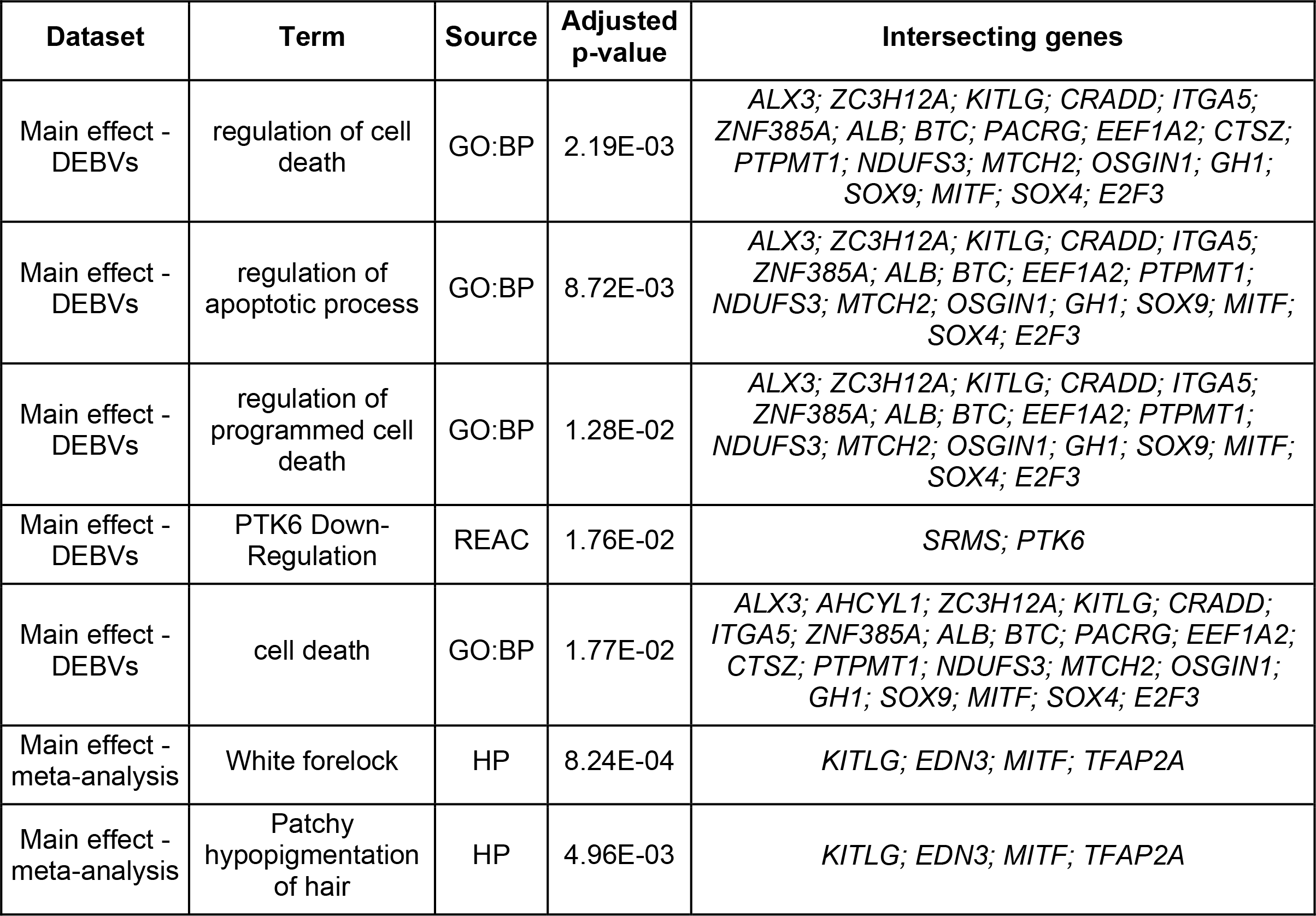
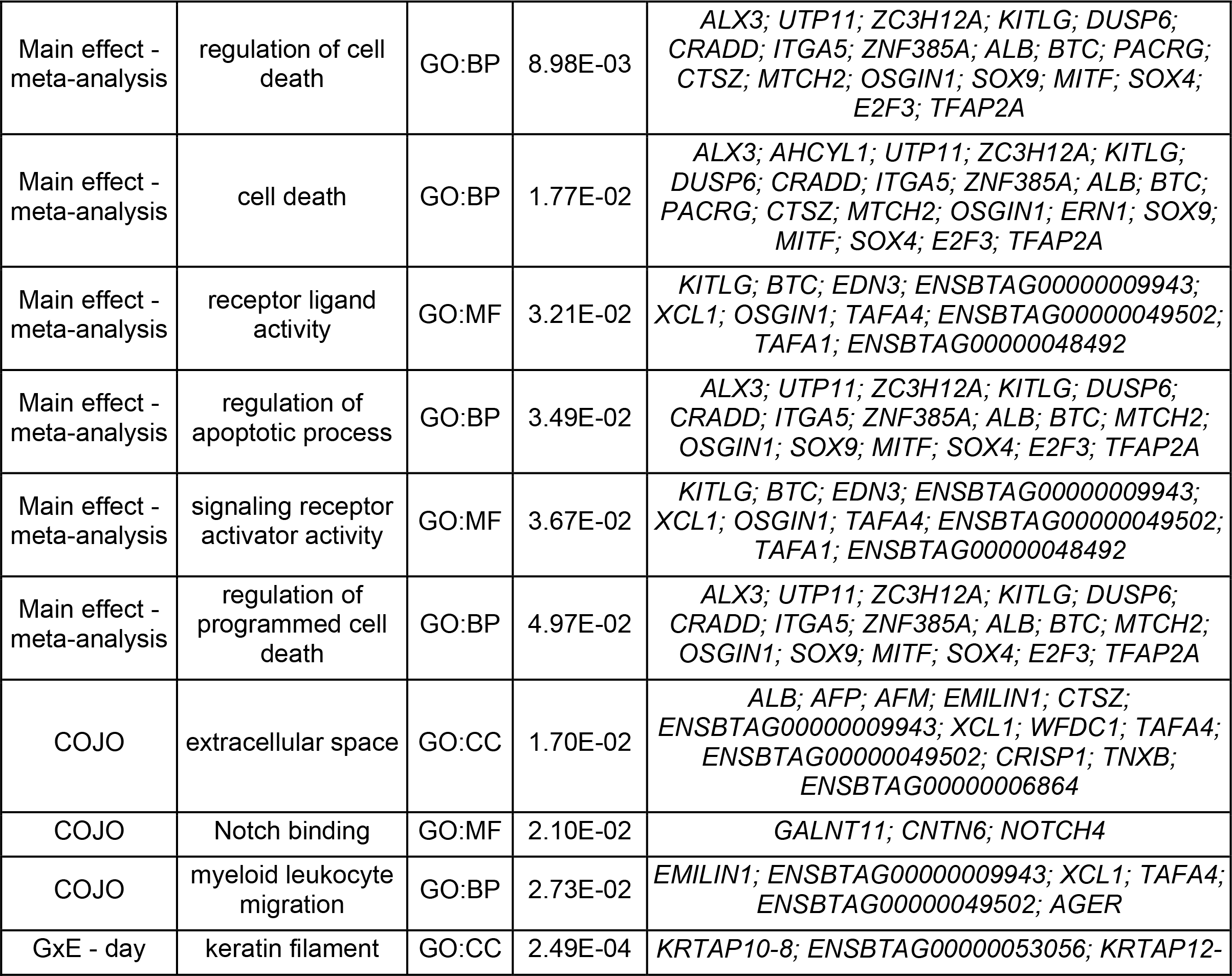
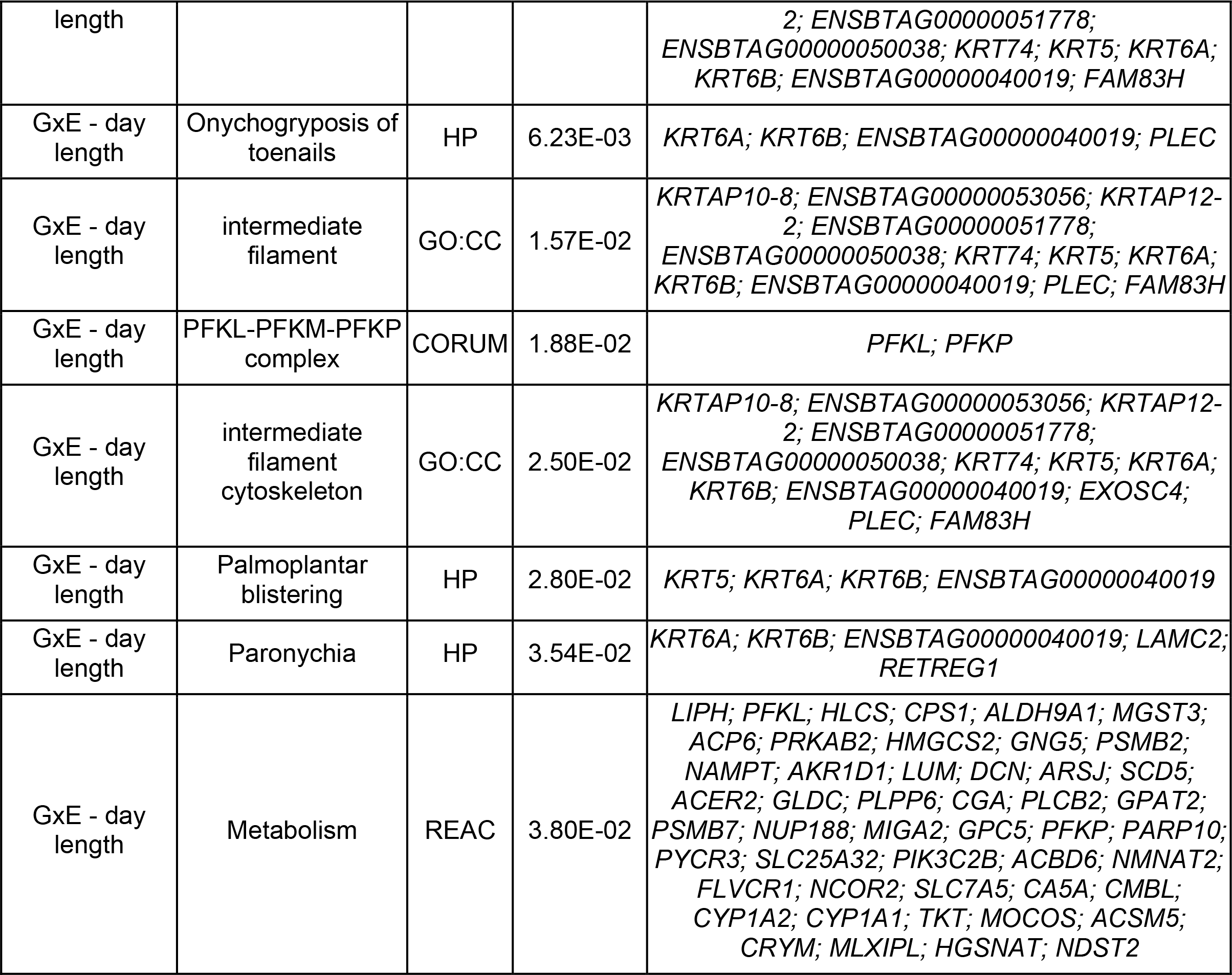
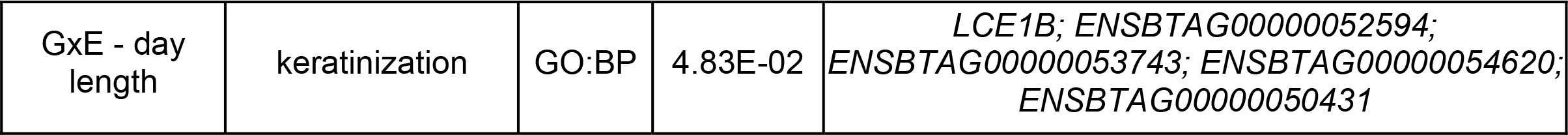
Gene set enrichment results. Considered data sources included GO biological processes, GO cellular components, GO molecular functions, KEGG pathways, Reactome pathways, CORUM pro, and disease phenotypes annotated by the Human Phenotype Ontology database. “Intersecting genes” are those genes in the set that were annotated to the term.

**Table S4.** QTL enrichment results. “N annotations” represent the number of Animal QTLdb annotations.

| Dataset | QTL name | Adjusted p-value | N annotations | N SNPs | SNPs |
| --- | --- | --- | --- | --- | --- |
| Main effect - DEBVs | White spotting | 8.35E-30 | 19 | 6 | 22:31,647,147; 22:32,120,539; 22:32,204,981; 22:32,208,077; 22:32,209,610; 22:32,979,644 |
| Main effect - DEBVs | Body weight (mature) | 7.28E-05 | 6 | 24 | 19:59,031,592; 19:59,042,109; 19:59,049,072; 19:59,058,798; 19:59,060,008; 19:59,062,908; 19:59,066,604; 19:59,069,136; 19:59,083,455; 19:59,084,200; 19:59,087,943; 19:59,091,866; 19:59,120,248; 19:59,133,155; 19:59,173,133; 19:59,179,679; 19:59,307,342; 19:59,309,672; 19:59,316,368; 19:59,327,548; 19:59,371,948; 19:59,379,387; 19:59,382,908; 19:59,393,728 |
| Main effect - DEBVs | Clinical mastitis | 2.11E-04 | 8 | 6 | 6:88,497,883; 13:57,010,672; 13:57,021,032; 13:57,029,389; 13:57,037,937; 13:57,744,564 |
| Main effect - DEBVs | Body weight (18 months) | 1.07E-02 | 2 | 18 | 5:18,834,893; 5:18,837,012; 5:18,837,973; 5:18,840,051; 5:18,845,572; 5:18,853,702; 5:18,861,017; 5:18,861,915; 5:18,863,193; 5:18,864,340; 5:18,865,142; 5:18,866,748; 5:18,876,734; 5:18,889,693; 5:18,924,994; 22:32,663,366; 22:32,718,597; 22:32,721,303 |
| Main effect - DEBVs | Sire conception rate | 2.03E-02 | 2 | 7 | 9:42,516,598; 13:57,939,566; 13:57,940,287; 13:57,958,615; 13:57,962,862; 13:57,971,981; 13:57,984,480 |
| Main effect - DEBVs | Eye area pigmentation | 3.82E-02 | 2 | 3 | 22:32,120,539; 22:32,663,366; 22:32,718,597 |
| Main effect - meta-analysis | White spotting | 6.34E-41 | 25 | 12 | 22:31,346,055; 22:31,601,001; 22:31,608,442; 22:31,647,147; 22:31,651,707; 22:31,653,105; 22:31,807,382; 22:32,120,539; 22:32,204,981; 22:32,208,077; 22:32,209,610; 22:32,979,644 |
| Main effect - meta-analysis | Body weight (mature) | 1.10E-05 | 7 | 28 | 5:22,351,623; 19:59,005,190; 19:59,009,634; 19:59,010,199; 19:59,011,273; 19:59,031,592; 19:59,042,109; 19:59,058,798; 19:59,062,908; 19:59,066,604; 19:59,069,136; 19:59,091,866; 19:59,103,428; 19:59,112,204; 19:59,120,248; 19:59,133,155; 19:59,167,705; 19:59,168,801; 19:59,173,133; 19:59,175,249; 19:59,179,679; 19:59,202,424; 19:59,307,342; 19:59,309,672; 19:59,316,368; 19:59,382,908; 19:59,391,271; 19:59,393,728 |
| Main effect - meta-analysis | Clinical mastitis | 5.37E-03 | 7 | 4 | 6:88,493,833; 6:88,497,883; 13:57,010,672; 13:57,037,937 |
| Main effect - meta-analysis | Body weight (18 months) | 2.45E-02 | 2 | 18 | 5:18,834,893; 5:18,837,012; 5:18,837,973; 5:18,840,051; 5:18,845,572; 5:18,853,702; 5:18,861,017; 5:18,861,915; 5:18,863,193; 5:18,864,340; 5:18,865,142; 5:18,866,748; 5:18,876,734; 5:18,889,693; 5:18,924,994; 22:32,663,366; 22:32,718,597; 22:32,721,303 |
| Main effect - meta-analysis | Hoof and leg disorders | 4.29E-02 | 2 | 4 | 22:31,608,442; 22:31,647,147; 22:31,651,707; 22:31,653,105 |
| COJO | Interval from first to last insemination | 2.83E-32 | 26 | 5 | 17:68,938,350; 17:69,096,404; 17:69,149,498; 17:69,155,480; 17:69,157,843 |
| COJO | Non-return rate | 3.43E-14 | 26 | 5 | 17:68,938,350; 17:69,096,404; 17:69,149,498; 17:69,155,480; 17:69,157,843 |
| COJO | Calf size | 3.07E-03 | 3 | 4 | 13:57,182,321; 13:57,201,678; 17:69,155,480; 17:69,157,843 |
| COJO | Bovine leukemia virus susceptibility | 1.74E-02 | 2 | 2 | 23:22,556,761; 23:27,294,315 |
| GxE - day length | Metabolic body weight | 6.12E-05 | 115 | 8 | 3:9,874,711; 14:1,073,049; 16:52,306,052; 20:4,609,314; 20:4,617,648; 20:4,633,738; 20:4,653,700; 20:4,655,604 |
| GxE - day length | Ketosis | 4.18E-03 | 21 | 13 | 3:32,993,574; 9:64,290,905; 13:52,556,307; 14:796,638; 14:1,073,049; 20:58,688,893; 20:58,742,109; 20:58,745,645; 20:58,748,682; 20:58,762,246; 20:58,792,088; 20:58,796,066; 20:59,139,410 |
| GxE - day length | Lifetime profit index | 8.47E-03 | 6 | 2 | 14:796,638; 14:1,073,049 |

Supplementary File 2. CSV file with list of genes within 50 kbp of the associated SNPs.

